# Autism is Associated with *in vivo* Changes in Gray Matter Neurite Architecture

**DOI:** 10.1101/2023.03.25.534208

**Authors:** Zachary P. Christensen, Edward G. Freedman, John J. Foxe

## Abstract

Postmortem investigations in autism have identified anomalies in neural cytoarchitecture across limbic, cerebellar, and neocortical networks. These anomalies include narrow cell mini-columns and variable neuron density. However, difficulty obtaining sufficient post-mortem samples has often prevented investigations from converging on reproducible measures. Recent advances in processing magnetic resonance diffusion weighted images (DWI) make *in vivo* characterization of neuronal cytoarchitecture a potential alternative to post-mortem studies. Using extensive DWI data from the Adolescent Brain Cognitive Development^sm^ (ABCD®) study 142 individuals with an Autism diagnosis were compared with 8971 controls using a restriction spectrum imaging (RSI) framework that characterized total neurite density (TND), its component restricted normalized directional diffusion (RND), and restricted normalized isotropic diffusion (RNI). A significant decrease in TND was observed in Autism in the right cerebellar cortex (β=-0.005, SE =0.0015, p=0.0267), with significant decreases in RNI and significant increases in RND found diffusely throughout posterior and anterior aspects of the brain, respectively. Furthermore, these regions remained significant in *post-hoc* analysis when the ASD sample was compared against a subset of 1404 individuals with other psychiatric conditions (pulled from the original 8971). These findings highlight the importance of characterizing neuron cytoarchitecture in Autism and the significance of their incorporation as physiological covariates in future studies.

**Lay abstract:** Children with autism have differences in neuron structure unique from the general population *and* populations with attention, anxiety, and depression disorders. Brain imaging data on over 11,000 children was acquired at ages 9 and 11 years-of-age. Estimates of neuron density were derived from brain imaging data using recently validated techniques and comparative groups were composed using parent reported diagnosis of autism and other common psychiatric disorders. Consistent macro-structural changes in brain have been difficult to replicate and micro-structural changes have been historically difficult to acquire with other methodologies. We identified regional differences in the density of neuron cell bodies, neuron branching, and total neuron density in those with a reported diagnosis of ASD. Findings were consistent when compared against those with other psychiatric disorders in post-hoc analysis. These findings demonstrate the viability and importance of investigating *in vivo* changes to neurons in those with autism to advance our current understanding of related physiology.

## 1 Introduction

Autism spectrum disorder (ASD) is a neurodevelopmental disorder characterized by core symptomatology of restricted interests, impaired social communication, and repetitive patterns of behavior (American Psychiatric Association & American Psychiatric Association, 2013). ASD symptomatology has been associated with differences in brain structure (Alemany et al., 2021). Children with ASD at 2-4 years of age have a larger cerebral volume compared to typically developing children (Courchesne, 2003; Lee et al., 2021). Previous efforts to explain this phenomenon have found that increased cortical surface area and increased cerebral volume co-occur in children with ASD (Hazlett et al., 2011). Rates of brain growth from 2 to 4 years of age are similar in children with ASD and typically developing children, suggesting that greater surface area in ASD is driven by an increase in the number or size of cortical gyri prior to two years of age (Hazlett et al., 2011). Greater cerebral volume and cortical surface area are not identified frequently after the age of four (Courchesne et al., 2011). Instead, children with autism have increased volume of the amygdala (Groen et al., 2010; Schumann, 2004; Schumann et al., 2009) and increased cortical thickness throughout the frontal lobe (Khundrakpam et al., 2017). Volumetric and cortical thickness measures appear to normalize as children progress into adolescence and adulthood (Courchesne, 2004; Courchesne et al., 2007; Khundrakpam et al., 2017).

The relationship between measures localized to specific brain regions and diffuse global brain measurements has been suggested to have an important role in the development of ASD. For example, cortical thickness and volumetric covariance measures have demonstrated utility in the development of diagnostic classifiers for ASD (Zheng et al., 2021). Diffusion weighted imaging (DWI) has consistently identified measures indicative of decreased fiber integrity throughout the brain of those with ASD (Conti et al., 2017; Groen, 2011; Libero et al., 2016). However, decreased fiber integrity is associated with many other neurodevelopmental and psychiatric diagnoses and may simply be a non-specific indicator of global perturbation (Benedetti et al., 2014; Stojanovski et al., 2019; Xu et al., 2022). Increased cortical hemispheric symmetry appears to be more specific to ASD but has proven difficult to replicate (Carper et al., 2016; Deng & Wang, 2021; Postema et al., 2019; Sha et al., 2021). Recent investigations have found that dysregulation of regional gene expression is associated with global cortical thickness (Romero-Garcia et al., 2019) and structural connectivity (Park et al., 2021). However, the relationship between different neuron populations throughout neurodevelopment in ASD remains poorly understood.

Current understanding of neural cytoarchitecture in ASD relies heavily on post-mortem studies that may reveal microscopic perturbations that are not reflected in macroscopic measures. For example, individuals with ASD have been found with a reduced number of neurons in the amygdala but no difference in overall volume of the amygdala (Schumann & Amaral, 2006). Early histological studies of ASD identified fewer cerebral Purkinje cells (Kemper & Bauman, 1998), smaller Purkinje cells in the cerebellum (Fatemi et al., 2002), patches of cortical disorganization (Stoner et al., 2014), and a greater number of narrow minicolumns in the frontal and temporal lobes (Buxhoeveden et al., 2006; M. F. Casanova et al., 2002, 2002; M. Casanova & Trippe, 2009). Many of these findings have not been clearly reproduced or have met with contradictory findings, such as evidence that minicolumns are in fact wider (McKavanagh et al., 2015). This may be due to the extremely small sample sizes of most post-mortem studies or comparison between individuals at different developmental stages with cytoarchitectural differences unrelated to ASD (Schumann & Nordahl, 2011).

Recent advances in DWI processing utilize multicompartment models to derive measures characterizing brain cytoarchitecture. Unlike traditional diffusion tensor imaging (DTI), multicompartment DWI measurements are not compromised by neuronal projections with non-linear paths or dense complex architectures, permitting better investigation of cortical cytoarchitecture and complex fiber tracts (Fukutomi et al., 2018, 2019; White et al., 2013; Zhang et al., 2012). The total neurite density (TND) is of particular interest and represents the fraction of tissue within a voxel that is composed of axons and dendrites. Compared to typical DTI measures, TND has higher accuracy and specificity in identifying histological differences. A recent investigation of 27 individuals with ASD found microstructural changes measured using TND had greater effect sizes than corresponding DTI measures (Carper et al., 2017). Furthermore, individuals with ASD have demonstrated a negative correlation between deficits in facial processing and TND in the ventral occipital complex, parietal areas, and forceps major (Yasuno et al., 2020). However, neurite density measures are sensitive to age (Genc et al., 2017; Qian et al., 2020), complicating discovery and validation of measures in neurodevelopmental disorders such as ASD.

The aim of the present study is to provide measures of ASD cytoarchitecture that overcome the uncertainty of previous findings due to small sample size and cytoarchitectural changes across development. This is accomplished through use of the Adolescent Brain and Cognitive Development^sm^ (ABCD®) study (Barch et al., 2018, 2021; Casey et al., 2018; Hagler et al., 2019), providing 11,873 participants between the ages of 9-10 years-old at baseline. Image acquisition and processing provide measures comparable to TND within regions of interest corresponding to a standard brain atlas. All derived measures are compared between those with ASD and those with no diagnosis of ASD (nASD). Finally, notable findings in ASD are compared against those with other psychiatric diagnoses (OPD), ensuring findings are specific to ASD.

## 2 Methods

### 2.1 Subjects

Data from the Adolescent Brain and Cognitive Development^sm^ (ABCD®) study were used (Casey et al., 2018). The ABCD study consists of 11,873 children recruited between ages 9-10-year-old at baseline. Participants were recruited from 22 sites across the US from public and private schools and selection was based on race and ethnicity, sex, socioeconomic status, and urbanicity, in an effort to match US population and demographics (Garavan et al., 2018). The ABCD study is currently in the process of following these individuals over the course of 10 years - https://abcdstudy.org/. The present study utilized completed timepoints with imaging data, baseline and the two-year follow-up, focusing on outcomes that were stable across both timepoints.

Individuals were excluded if they were not able to complete all administered tasks or complete an MRI scan. This excluded individuals who lacked fluency in English or uncorrectable sensory deficits (e.g., legal blindness). Participants were also excluded if they had a psychiatric diagnosis that prevented attendance at regular classes in school, neurological issues (e.g., brain tumor, previous head injury with loss of consciousness > 30 minutes), or significant perinatal medical issues (e.g., gestational age < 28 weeks, birthweight <1.2 kg, complications resulting in >1 month of hospitalization following birth) (Garavan et al., 2018). Given the focus of the current investigation, individuals that were missing pertinent imaging data were excluded (see *Neuroimaging Measures* section).

### 2.2 Diagnostic Groups and Cognitive Assessments

The ABCD cohort was subdivided into those with a parent reported diagnosis of ASD and those without a parent reported diagnosis of ASD (nASD). The brief social responsiveness scale (SRS) was used to overcome the absence of formal testing or clinical records to confirm a diagnosis of ASD. A T-score of above 60 on the SRS has demonstrated good reliability and sensitivity for identifying ASD across various psychopathologies (Moul et al., 2015). Participants who had a parent reported diagnosis of ASD and scored below this threshold on the SRS were removed from subsequent analysis to improve validity of diagnostic categories. An additional subgroup was identified from the nASD sample, and characterized as those with other psychiatric diagnoses (OPD), such as anxiety, depression, and ADHD. Comparisons between ASD and the OPD group were used to ensure findings were associated with pathology specific to ASD and not comorbidities in the OPD group in *post-hoc* analyses.

Given the association of ASD with a variety of behavioral outcomes (social function (Sugranyes et al., 2011), emotional regulation (Weiss, 2014), anxiety (Hessl et al., 2020), and attentional difficulties (Rodriguez‐Seijas et al., 2019)) measures from the Child Behavioral Checklist (CBCL) were used. The CBCL utilizes parent reports of behavior across a number of items to produce subscales of problematic behavioral severity. This has previously proven useful in analyzing behavioral outcomes for large pediatric cohorts (Barch et al., 2018; Michelini et al., 2019; Thompson et al., 2019; Tiemeier et al., 2012). Subscales estimating severity of problematic behaviors related to anxiety-depression, social functioning, attention, and somatization were used in the present study.

### 2.3 Neuroimaging Methods

Diffusion weighted images (DWI) underwent standard preprocessing as previously described in detail (this included corrections for eddy currents, motion, field inhomogeneities as well as other potential sources of artifact) (Casey et al., 2018). In brief, diffusion weighted images were fit to a restriction spectrum imaging (RSI) model (White et al., 2013). RSI provides multicompartmental measures of diffusion unique to intracellular (restricted) and extracellular (hindered) compartments of brain tissue. TND represents the summed estimates of restricted normalized directional diffusion (RND; corresponding to axonal structures) and restricted normalized isotropic diffusion (RDI; corresponding to the neuron body). These measures were assessed in 87 gray matter regions corresponding to cortical and subcortical regions found in the Desikan-Killiany brain atlas (Klein & Tourville, 2012). All measures were pulled from the latest curated ABCD data release.

### 2.4 Statistical Analysis

Each TND region and closeness centrality measure was evaluated as a response in a mixed effect linear model, using sex, highest parent education, collection site, reported use of medications for ADHD (methylphenidate, Ritalin, Adderall, etc.), and composite intelligence score, to compose the null model. Each of these models was compared to the null model via likelihood ratio tests. Primary analysis focused on measures pooled across baseline and two-year follow-up (ages 9-10 and 11-12, respectively), using subject ID as an additional control for measures attributable to a single individual across both timepoints. Post-hoc analysis was used to confirm significant findings were stable across baseline and two-year follow-up separately. Neurophysiological measures that were found to be significant were investigated as interacting terms with diagnostic information, using CBCL measures as outcomes. The resulting ANCOVA allowed us to informally compose an indirect effect measure, further describing how ASD influences physiology which in turn results in behavioral outcomes. Benjamini-Hochberg correction was used to correct for multiple comparisons (Benjamini & Hochberg, 1995). Code used to execute the analysis may be found at https://github.com/CognitiveNeuroLab/AutismNeuriteDensity upon publication.

## 3 Results

### 3.1 Sample Description

Analysis at both timepoints included 21,385 observations (11,210 at baseline and 10,175 at two-year follow-up). 310 of these observations were attributable to those in the ASD group (164 at baseline and 146 at two-year follow-up). 2,916 were attributable to those in the OPD group (1,538 at baseline and 1,378 at follow-up).

### 3.2 Association Between ASD and Behavioral Measures

Problematic behaviors were increased in the ASD group across all scales (anxiety-depression, attention, social somatic; see Table 1). These findings were consistent when examining only baseline observations (see Table 2) and follow-up observations (see Table 3). The ASD group also had increased problematic behaviors compared directly against the OPD group across all scales. However, only findings related to anxiety-depression and social behaviors were statistically significant (see Table 4).

**Table 1.** All time points

| **CBCL Scale** | **$\beta$ (SE)** | **p-value** |
| --- | --- | --- |
| Anxiety-depression | 0.841 (0.061) | <0.001 |
| Attention | 1.15 (0.059) | <0.001 |
| Social | 1.32 (0.061) | <0.001 |
| Somatic | 0.49 (0.061) | <0.001 |

**Table 2.** Baseline

| **CBCL Scale** | **$\beta$ (SE)** | **p-value** |
| --- | --- | --- |
| Anxiety-depression | 0.78 (0.080) | <0.001 |
| Attention | 1.21 (0.079) | <0.001 |
| Social | 1.38 (0.081) | <0.001 |
| Somatic | 0.43 (0.019) | <0.001 |

**Table 3.**
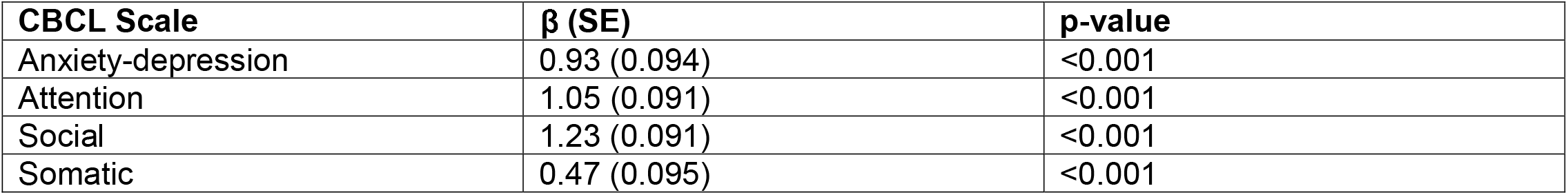
Two-year follow-up

| **CBCL Scale** | **$\beta$ (SE)** | **p-value** |
| --- | --- | --- |
| Anxiety-depression | 0.93 (0.094) | <0.001 |
| Attention | 1.05 (0.091) | <0.001 |
| Social | 1.23 (0.091) | <0.001 |
| Somatic | 0.47 (0.095) | <0.001 |

**Table 4.** All time points OPD vs ASD

| **CBCL Scale** | **$\beta$ (SE)** | **p-value** |
| --- | --- | --- |
| Anxiety-depression | 0.287 (0.087) | 0.0009 |
| Attention | 0.129 (0.079) | 0.103 |
| Social | 0.688 (0.091) | <0.001 |
| Somatic | 0.116 (0.081) | 0.149 |

### 3.3 Association Between ASD and MRI Measures

#### 3.3.1 Association Between TND and ASD

Total neurite density, combining measures from both baseline and two-year follow-up, was lower in occipital, parietal, midbrain, and hindbrain regions in the ASD group. Neurite density was greater in the frontal and temporal lobes of those with ASD. However, only the right cerebellar cortex was found to be significantly lower following correction for multiple comparisons (β=-0.005, SE =0.0015, p=0.0267; see Figure 1). Post-hoc analysis found the relationship at the right cerebellar cortex was consistent when evaluating baseline measures in isolation (β =-0.005, SE=0.0019, p=0.010), follow-up measures in isolation (β=-0.005, SE=0.002, p=0.013), and when comparing the ASD group only to the OPD group (β =-0.005, SE=0.002, p=0.003).

**Figure 1.**
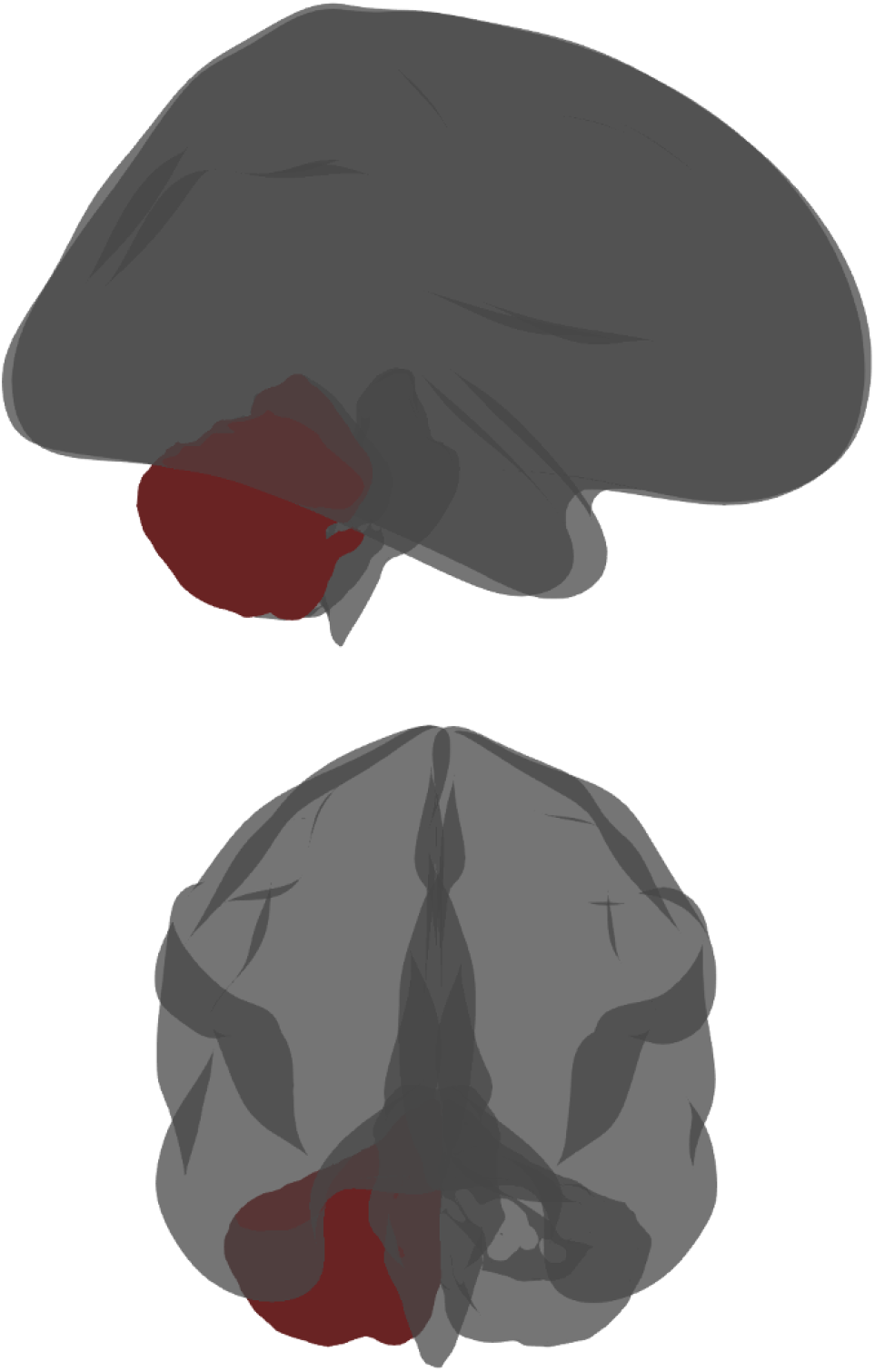

#### 3.3.2 Association Between RND and ASD

A pattern of increased restricted normalized directional (RND) diffusion pooled across baseline and two-year follow-up was found in the ASD group throughout the temporal lobe, frontal lobe, and cingulate gyrus. This was in contrast with a strong negative relationship between autism and RND in the left and right ventral diencephalon and brainstem (see Table 5 and Figure 2). Isolated analysis of baseline data found a pattern of increased RND in the ASD group at the left and right caudal anterior cingulate (β=0.283, SE=0.078, p=0.014 and β=0.275, SE=0.078, p=0.001), decreased RND at the left and right ventral diencephalon (β=-0.217, SE=0.068, p=0.029 and Β=-0.248, SE=0.069, p=0.014), and decreased RND at the brainstem (β=-0.201, SE=0.064, p=0.002). Isolated analysis of follow-up data found a pattern of increased RND in the ASD group at the left and right putamen (β =0.254, SE=0.062, p=0.002 and Β=0.259, SE=0.062, p=0.002), the left caudate (β =0.245, SE=0.062, p=0.002), the left amygdala (β=0.231, SE=0.078, p=0.041), the right insula (β=0.188, SE=0.055, p=0.010), and the right pars opercularis (β=0.251, SE=0.073, p=0.010)

**Table 5.** Statistically significant RND findings when analyzing all timepoints.

| ROI | Hemisphere | $\beta \pm SE$ | p-value |
| --- | --- | --- | --- |
| lateral orbitofrontal | left | 0.146 ± 0.053 | 0.02578364 |
| pars orbitalis | left | 0.172 ± 0.061 | 0.02248895 |
| bankssts | left | 0.17 ± 0.058 | 0.01844477 |
| entorhinal | left | 0.162 ± 0.051 | 0.0124094 |
| caudal anterior cingulate | left | 0.199 ± 0.059 | 0.00841803 |
| caudate | left | 0.218 ± 0.05 | 0.00055636 |
| putamen | left | 0.151 ± 0.048 | 0.0124094 |
| ventral diencephalon | left | -0.19 ± 0.052 | 0.00560059 |
| lateral orbitofrontal | right | 0.175 ± 0.054 | 0.01109968 |
| pars opercularis | right | 0.185 ± 0.057 | 0.01109968 |
| pars orbitalis | right | 0.213 ± 0.061 | 0.00841803 |
| caudal middle frontal | right | 0.158 ± 0.058 | 0.02578364 |
| supramarginal | right | 0.182 ± 0.059 | 0.01583416 |
| inferior temporal | right | 0.157 ± 0.054 | 0.02091586 |
| middle temporal | right | 0.148 ± 0.056 | 0.03303631 |
| bankssts | right | 0.17 ± 0.056 | 0.01590182 |
| caudal anterior cingulate | right | 0.176 ± 0.058 | 0.01590182 |
| insula | right | 0.135 ± 0.047 | 0.02166713 |
| caudate | right | 0.146 ± 0.051 | 0.02166713 |
| putamen | right | 0.168 ± 0.049 | 0.00841803 |
| accumbens | right | 0.124 ± 0.048 | 0.0370369 |
| ventral diencephalon | right | -0.237 ± 0.053 | 0.00055636 |
| brain stem | | -0.179 ± 0.049 | 0.00560059 |

**Figure 2.**
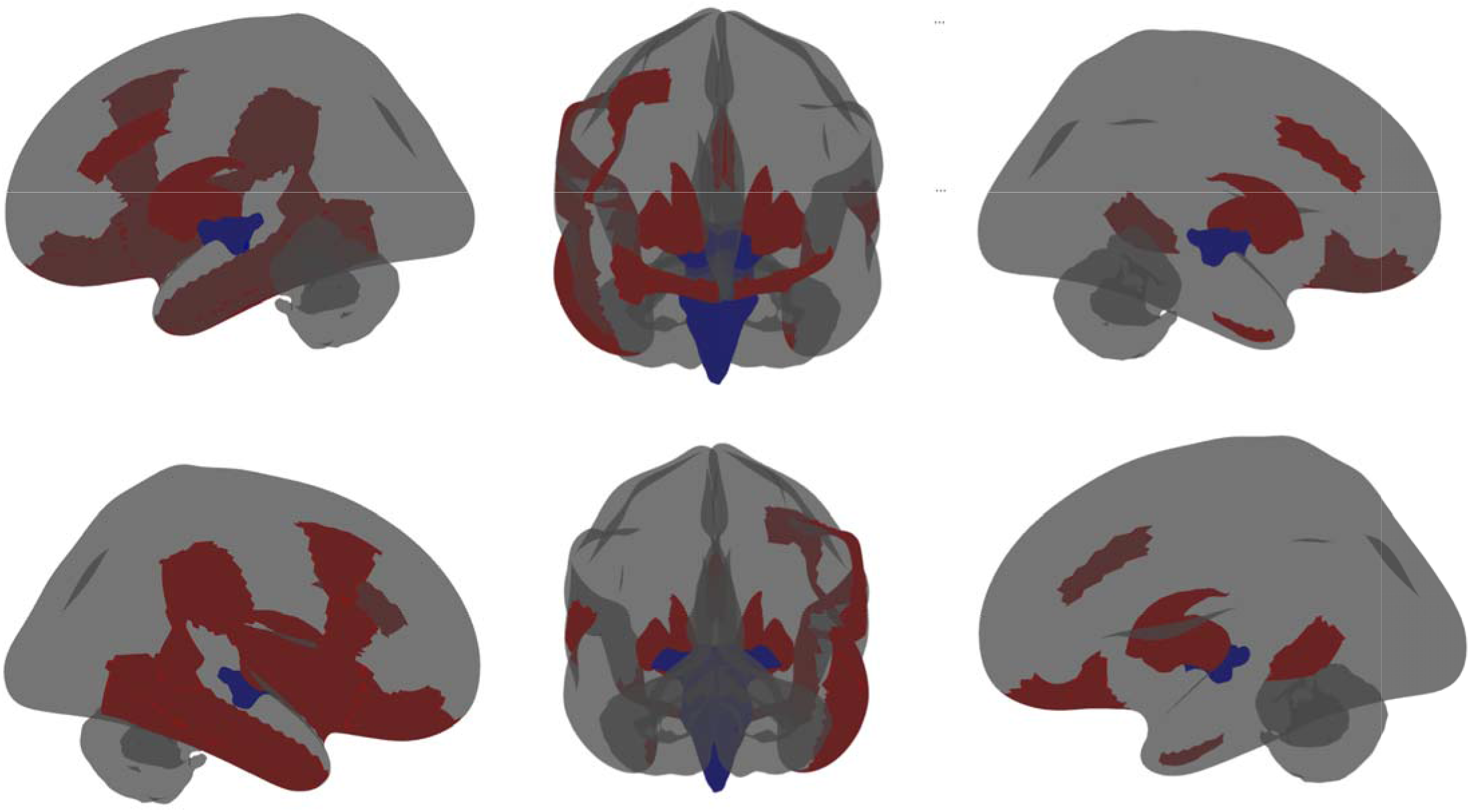

Comparing the ASD and OPD groups also identified decreased RND in the ASD group at the left and right ventral diencephalon (β=-0.194, SE=0.053, p=0.007 and β=-0.235, SE=0.055, p=0.002) and brain stem (Β=- 0.188, SE=0.052, p=0.007). Isolated analysis of baseline data found decreased RND in the ASD group at the left and right ventral diencephalon (β=-0.217, SE=0.067, p=0.049 and β=-0.248, SE=0.070, p=0.032). Examination of follow-up data alone found comparable findings as those at follow-up when analyzing the entire population (e.g., increased RND in the left and right putamen and right caudate), but none remained statistically significant after correction for multiple comparisons.

#### 3.3.3 Association Between RNI and ASD

A pattern of decreased RNI was found in the ASD group throughout the parietal lobe, occipital lobe, midbrain, and hindbrain (see Table 6). RNI was greater at the right and left amygdala in the ASD group (β=-0.006, SE=0.002, p=0.003; Β=0.003, SE=0.001, p=0.011; see Figure 3). Post-hoc analysis found that most regions remained significant when comparing ASD against the OPD group (see Table 7). Post-hoc analysis of baseline data found consistently significant findings at the left paracentral cortex (B=-0.005, SE = 0.0017, p=0.035), left postcentral cortex (B=-0.003, SE=0.008, p=0.035), left cerebellar cortex (B=-0.005, SE= 0.002, p=0.035), right paracentral cortex (B=-0.005, SE=0.0017, p=0.009), right superior temporal cortex (B=-0.002, SE=0.013, p=0.041), right lateral occipital cortex (B=-0.003, SE=0.010, p=0.035), and right cerebellar cortex (B=-.006, SE=0.001, p=0.010). Post-hoc analysis of follow-up data found consistently significant findings at the left paracentral cortex (B=-0.005, SE=0.002, p=0.041), left transverse temporal region (B=-0.005, SE=0.001, p=0.041), left superior temporal cortex (B=-0.004, SE=0.001, p=0.041), left pericalcarine region (B=-0.005, SE=0.002, p=0.041), left cuneus (B=-0.005, SE=0.002, p=0.041), left amygdala (B=0.006, SE=0.002, p=0.041), right cuneus (B=-0.005, SE=0.002, p=0.041), right amygdala (B=0.004, SE=0.002, p=0.041), and right cerebellar cortex (B=-0.006, SE=0.002, p=0.044).

**Table 6.** Statistically significant RNI findings when analyzing all timepoints.

| ROI | Hemisphere | $\beta \pm SE$ | p-value |
| --- | --- | --- | --- |
| paracentral | left | -0.00508 0.0014 | 0.01219271 |
| postcentral | left | -0.0035 ± 0.0011 | 0.02219444 |
| transverse temporal | left | -0.00361 ± 0.0013 | 0.0292521 |
| superior temporal | left | -0.00262 ± 0.00087 | 0.02607908 |
| lingual | left | -0.00283 ± 0.0011 | 0.04800862 |
| pericalcarine | left | -0.00371 ± 0.0013 | 0.0292521 |
| cuneus | left | -0.00338 ± 0.0013 | 0.04800862 |
| isthmus cingulate | left | -0.00265 ± 0.001 | 0.04800862 |
| amygdala | left | 0.0034 ± 0.0013 | 0.04800862 |
| cerebellum | left | -0.00495 ± 0.0016 | 0.02309396 |
| paracentral | right | -0.00486 ± 0.0014 | 0.01219271 |
| postcentral | right | -0.00305 ± 0.0011 | 0.0292521 |
| superior parietal | right | -0.00314 ± 0.0011 | 0.0292521 |
| inferior parietal | right | -0.00246 ± 0.00098 | 0.04800862 |
| superior temporal | right | -0.00279 ± 0.00086 | 0.02219444 |
| lingual | right | -0.00339 ± 0.0011 | 0.02309396 |
| lateral occipital | right | -0.00374 ± 0.0013 | 0.02646444 |
| pericalcarine | right | -0.00347 ± 0.0013 | 0.03022697 |
| cuneus | right | -0.00391 ± 0.0013 | 0.02607908 |
| isthmus cingulate | right | -0.00274 ± 0.001 | 0.03693149 |
| amygdala | right | 0.00362 ± 0.0013 | 0.0292521 |
| cerebellum | right | -0.00639 ± 0.0016 | 0.00380574 |

**Table 7.** Statistically significant RNI findings when comparing ASD against only OPD aftering correcting for multiple comparisons.

| ROI | Hemisphere | $\beta \pm SE$ | p-value |
| --- | --- | --- | --- |
| paracentral | left | -0.004 ± 0.002 | 0.025 |
| postcentral | left | -0.003 ± 0.001 | 0.032 |
| superior temporal | left | -0.002 ± 0.001 | 0.032 |
| pericalcarine | left | -0.003 ± 0.001 | 0.047 |
| cerebellum | left | -0.003 ± 0.002 | 0.032 |
| paracentral | right | -0.003 ± 0.002 | 0.025 |
| postcentral | right | -0.003 ± 0.001 | 0.032 |
| superior temporal | right | -0.002 ± 0.001 | 0.032 |
| lingual | right | -0.003 ± 0.001 | 0.032 |
| lateral occipital | right | -0.003 ± 0.001 | 0.032 |
| pericalcarine | right | -0.003 ± 0.001 | 0.032 |
| cuneus | right | -0.003 ± 0.001 | 0.032 |
| Isthmus cingulate | right | -0.003 ± 0.001 | 0.032 |
| amygdala | right | 0.004 ± 0.001 | 0.025 |
| cerebellum | right | -0.003 ± 0.001 | 0.025 |

**Figure 3.**
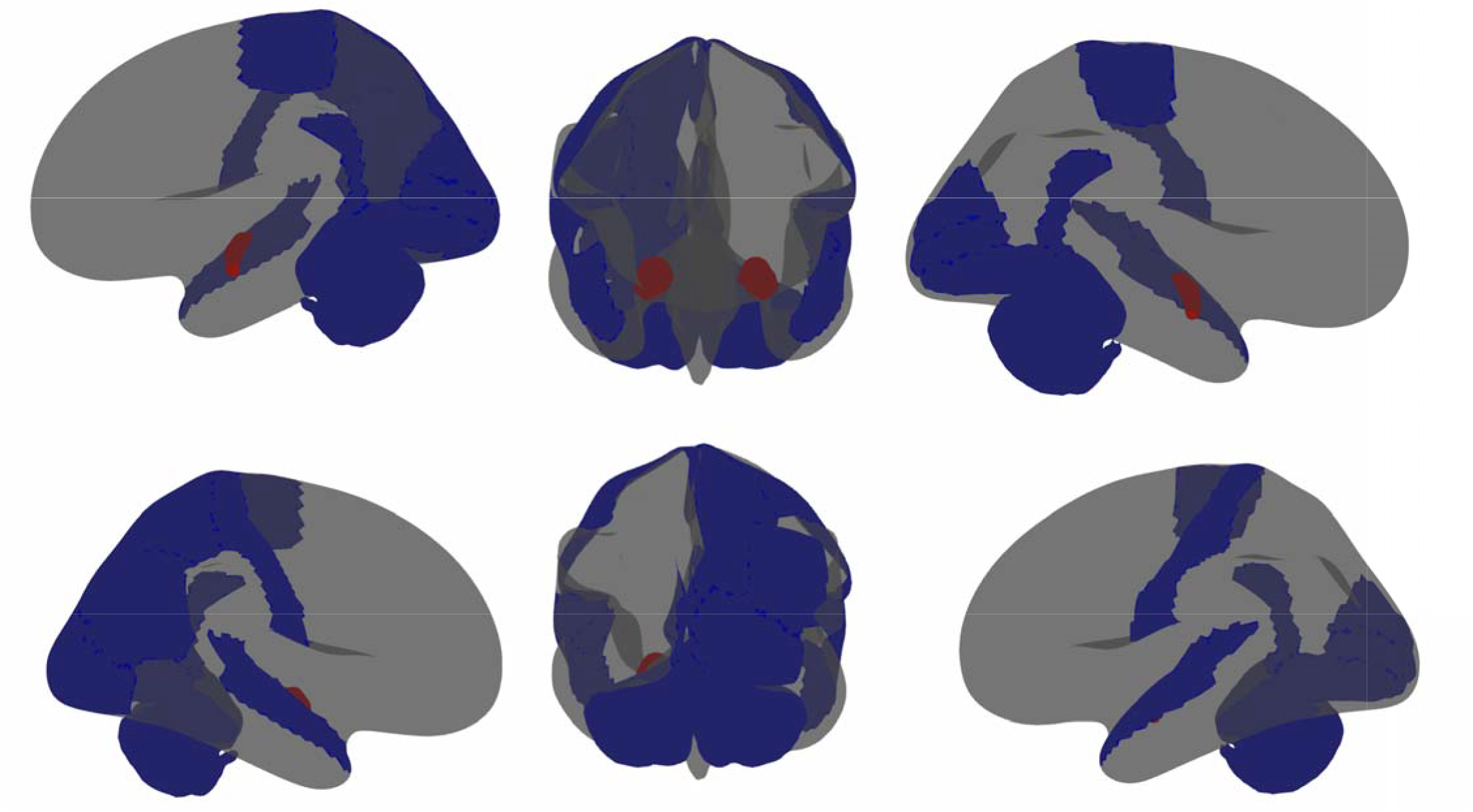

#### 3.3.4 Association Between FA and ASD

Post-hoc examination of fractional anisotropy values provided comparable measures using a DTI model. A pattern of increased FA in the frontal lobe and cingulate gyrus was found in the ASD group and a pattern of decreased FA was found in the left and right ventral diencephalon and brainstem in the ASD group (see Table 8). These findings were similar to those of the RND analysis but had a general decrease in effect size, resulting in the absence of several ROIs identified in the RND analysis (such as those in the temporal lobe).

**Table 8.** Statistically significant FA findings when analyzing all timepoints. (STS=superior temporal sulcus)

| ROI | Hemisphere | $\beta \pm SE$ | p-value |
| --- | --- | --- | --- |
| pars orbitalis | left | 0.167 ± 0.06 | 0.04403367 |
| caudal anterior cingulate | left | 0.199 ± 0.057 | 0.01041919 |
| caudate | left | 0.247 ± 0.052 | 0.00010585 |
| ventral diencephalon | left | -0.156 ± 0.041 | 0.00340059 |
| lateral orbitofrontal | right | 0.141 ± 0.052 | 0.04561598 |
| pars orbitalis | right | 0.166 ± 0.06 | 0.04403367 |
| caudal middle frontal | right | 0.164 ± 0.057 | 0.04392397 |
| supramarginal | right | 0.172 ± 0.058 | 0.04236129 |
| Banks STS | right | 0.163 ± 0.057 | 0.04392397 |
| caudal anterior cingulate | right | 0.161 ± 0.057 | 0.04392397 |
| ventral diencephalon | right | -0.195 ± 0.04 | 0.00010585 |
| brain stem |  | -0.146 ± 0.042 | 0.01041919 |

Isolated analysis of baseline data found a pattern of increased FA in the ASD group at the left and right caudal anterior cingulate (β=0.279, SE=0.078, p=0.010 and β=0.251, SE=0.078, p=0.026) and decreased FA at the left and right ventral diencephalon (Β=-0.184, SE=0.052, p=0.010 and Β=-0.202, SE=0.052, p=0.008). Analysis of baseline data also identified a pattern of decreased FA in the brain stem that was not statistically significant. Isolated analysis of follow-up data found increased FA in the ASD group at the left and right caudate (Β=0.315, SE=0.061, p<0.001 and Β=0.189, SE=0.060, p=0.003), the left and right putamen (β=0.279, SE=0.064, p=0.001 and Β=0.231, SE=0.060, p=0.003), the banks of the right superior temporal sulcus (β=0.244, SE=0.074, p=0.023), and decreased FA at the right ventral diencephalon (Β=-0.183, SE=0.059, p=0.027).

Examination of ASD against just the OPD group identified decreased FA in the ASD group at the left and right ventral diencephalon (β=-0.157, SE=0.042, p=0.004 and β =-0.183, SE=0.042, p=0.001) and brain stem (β=- 0.167, SE=0.045, p=0.0002). Isolated analysis of baseline data found decreased FA in the ASD group at the left and right ventral diencephalon (β=-0.188, SE=0.052, p=0.013 and β=-0.201, SE=0.053, p=0.012). Isolated analysis of follow-up data found increased FA in the ASD group at the left caudate (β =0.282, SE=0.077, p=0.0002).

#### 3.3.5 Association Between TND and Behavioral Outcomes

Post-hoc analysis between behavioral outcomes and TND at the right cerebellar cortex demonstrated that decreases in TND were associated with increased behavioral problems. This association was weak for measures of problematic outcomes on social (B=-1.951, SE=2.633, p=0.459), anxiety-depression (B=-4.864, SE=2.64, p=0.066), and attention (B=-0.585, SE=2.581, p=0.821). However, the relationship between TND at the right cerebellar cortex and somatization was strong (B=-7.1887, SE=2.67058, p=0.007; see Figure 4). This relationship remained strong when comparing the ASD group against just the OPD group (B=-8.371, SE=3.477, p=0.016) and when only evaluating measures at baseline (B=-7.702, SE=3.274, p=0.019). This association was much weaker when evaluating measures only at follow-up (B=-6.272, SE=4.675, p=0.180).

**Figure 4.**
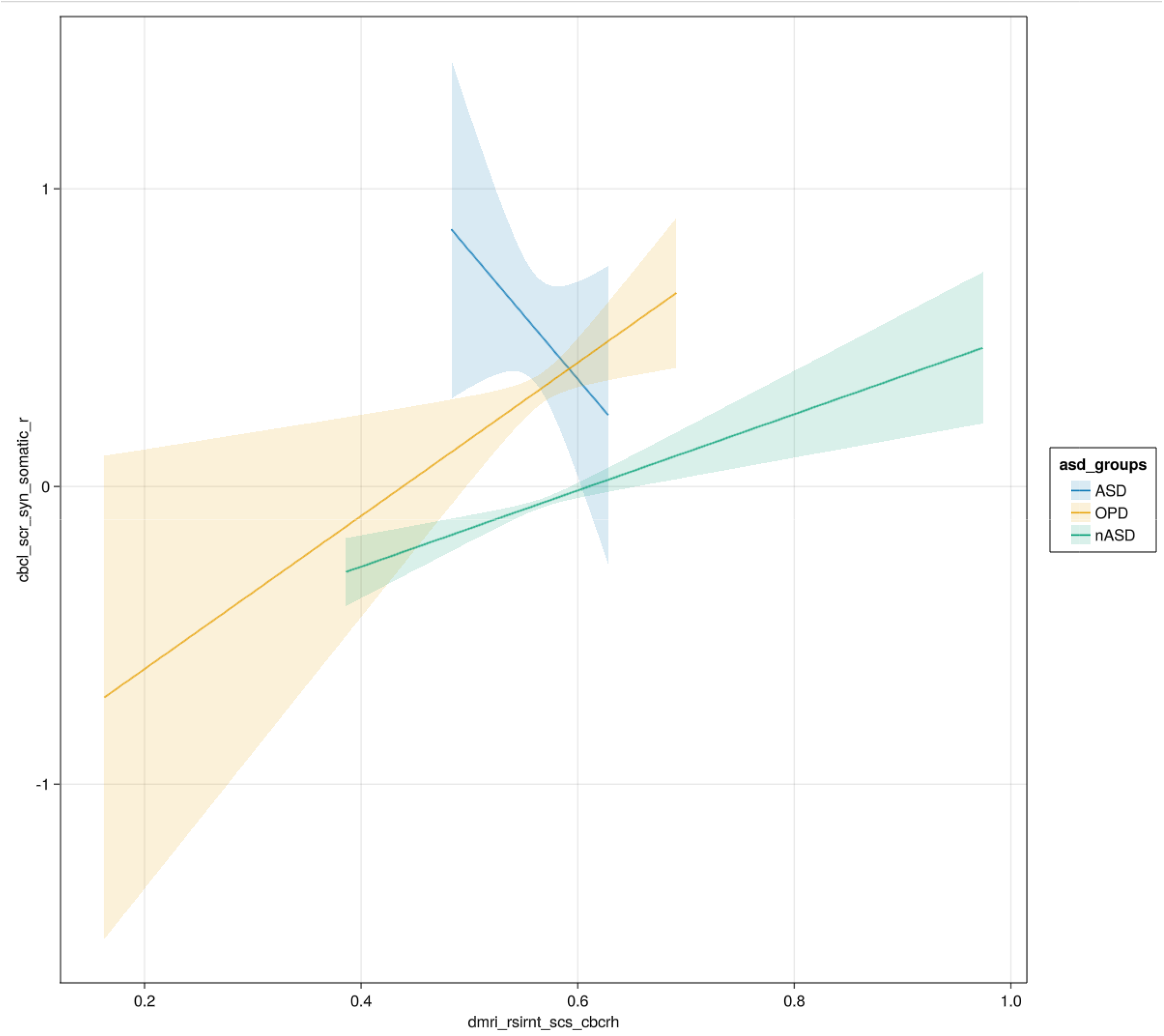
Correlation between TND and somatization for ASD, OPD, and nASD separately.

## 4 Discussion

The present study is the largest analysis of gray matter cytoarchitectural microstructure in ASD using a multi-compartmental diffusion model driven by multi-shell MRI. Total neurite density (TND) in the right cerebellar cortex was found to be lower in the ASD group. However, splitting TND into restricted normalized directional (RND) and restricted normalized isotropic (RNI) diffusion revealed unique regional changes associated with a diagnosis of ASD. Measures of RND in the ASD group were greater in frontotemporal regions and lower in the ventral diencephalon and brain stem. In contrast, measures of RNI were lower through parietooccipital regions and higher at the amygdala. Post-hoc analysis supported the consistency of these findings when isolating baseline and follow-up time points and when comparing ASD to other common psychiatric disorders, suggesting that these findings are truly specific to ASD. Finally, TND at the cerebellum had a significant association with behavioral outcomes related to somatic complaints.

These findings indicate that more intricate measures of *in vivo* cytoarchitecture in ASD allow identification of novel findings, otherwise obscured by macroscopic measurements or more traditional summary statistics of cytoarchitecture.

### 4.1 Neurite Density

Previous evaluation of RSI-based neurite density measures struggled to find significant group effects in ASD (Carper et al., 2017). However, this may be attributable to a smaller sample size and distributed age range, given the present study found decreased TND in the right cerebellar cortex of the ASD group. Although other regions had lower TND in the ASD group (such as the left cerebellar cortex and brain stem), none survived correction for multiple comparisons. Globally reduced TND is to be expected based on previous evaluation of RSI measures in ASD (Carper et al., 2017).

Separating TND into its components (RND and RNI) provided unique insights into the characteristics of neurons throughout the brain. RNI is assumed to correspond to density of neuron cell bodies and RND is assumed to correspond to dendritic and axonal structures (Palmer et al., 2022; White et al., 2013). Therefore, the present study’s findings may be interpreted as showing that those with ASD have decreased neuron cell body density in the parietal lobe, occipital lobe, midbrain, hindbrain, and decreased density of neuron branching in the ventral diencephalon and cerebellum. In contrast, those with ASD also show increased density of neuron branching in fronto-temporal, caudate, insula, and putamen regions and increased neuron cell body density in the amygdala.

### 4.2 ASD vs OPD

ASD has a rich history of comorbidities and associative analysis with other psychiatric disorders and behavioral anomalies (e.g. anxiety, ADHD, somatization) (Hessl et al., 2020; Rodriguez□Seijas et al., 2019; Sugranyes et al., 2011; Weiss, 2014). This may be useful in furthering awareness of needs and difficulties in those with ASD, but it also obscures the distinctiveness of biological associations to ASD. For example, ADHD and ASD demonstrate comorbidity and an association with structural abnormalities in regions related to executive functioning (Di Martino et al., 2013; Kangarani□Farahani et al., 2022). These similarities could represent a common pathogenesis, a convergence of physiological changes, or something else entirely.

In the present study the OPD group was pulled out of the overall non-ASD group for separate post-hoc comparisons, containing individuals with a reported history of ADHD, anxiety, depression, and other common psychiatric disorders. Although the ASD group had significantly higher reported problematic behaviors related to anxiety-depression, social functioning, somatization, and attention; only anxiety-depression and social behaviors demonstrated significant differences when compared to the OPD group. Furthermore, post-hoc analysis of all significant cytoarchitectural findings remained strongly associated when independently compared between ASD and OPD groups. The unique association between cytoarchitectural measures and behavioral outcomes in ASD may be due to the unique developmental origins of ASD from other disorders.

Testing this hypothesis and the mechanism by which this occurs remains an important area for future research to explore.

### 4.3 Longitudinal Changes

ASD has historically been associated with macro-structural brain changes overtime (Khundrakpam et al., 2017; Lange et al., 2015; Lee et al., 2021). Although these changes have not been easily reproduced, they constitute the only available structural measure monitored over time, due to obvious limitations in post-mortem studies.

The present study measures cytoarchitecture at two time-points per-subject across a two-year span (9 and 11 years-of-age). Post-hoc analysis demonstrated that evaluation of each time-point separately yielded similar findings. This may indicate that cytoarchitecture remains consistent in the presence of ongoing macro-structural changes. However, this represents a limited time frame across the developmental timeline pertinent to ASD and isn’t entirely consistent with previous claims. For example, Schuman and Amaral found reduced neurons in amygdala but no change in volume (Schumann & Amaral, 2006). Therefore, these findings should be interpreted only as an initial step in a much more exhaustive evaluation of longitudinal cytoarchitectural changes in ASD.

### 4.4 Relation to Behavior Findings

Significant TND findings were associated with behavioral outcomes to evaluate the potential of the present study’s findings for clinical use. This revealed a uniquely strong negative correlation between ASD and somatization that held up when also isolating comparison of those with ASD to those with OPD and when only using measures at baseline. It is possible that the decreased correlation between TND and somatization at follow-up is due to the smaller sample size at follow-up, as evidenced by the increased standard error. It is also possible that the shift in findings is due to ongoing developmental changes in neural cytoarchitecture throughout the two years between baseline and follow-up. As the current study has endeavored to establish the importance of neural cytoarchitecture and its potential relevance to behavioral outcomes in ASD, further investigation is necessary to explore the mechanism by which these two are connected.

### 4.5 Comparison to Other Measures

TND represents the most traditional measure in the present study. In previous literature, it has been compared to the neurite density index (NDI) used in the NODDI multicompartment model (Palmer et al., 2022), which has been in turn related to FA from traditional DTI models (Carper et al., 2017). These comparisons have predominantly been applied to cytoarchitecture in white matter where the DTI model is most accurate. A few studies have directly compared NDI to FA in gray matter, demonstrating similar findings (but with greater effect sizes using NDI). Comparisons to RSI measures and traditional DTI models aren’t as well studied in the literature but show a similar trend (Beck et al., 2021). Interestingly, the present study found FA measures to be more consistent with RND measures.

The association between FA and RND is not entirely unexpected since FA and RND both represent the directional or non-isotropic part of the diffusion signal. However, this does pose an issue when comparing to previous literature where neurite density has been conceptually aligned with FA in many instances. For example, previous literature claimed that TND was not consistent with histological findings related to prefrontal axons in ASD (specifically the orbitofrontal cortex) (Zikopoulos & Barbas, 2010). However, the present study identified numerous changes in the prefrontal cortex, including the pars orbitalis and lateral orbitofrontal cortex.

### 4.5 Limitations

The present study has some limitations, particularly those inherent to use of diagnostic information currently available in the ABCD dataset. ASD is typically diagnosed using the Autism Diagnostic Observation Schedule (ADOS) (Lord et al., 2000). The ABCD study does not provide any formal diagnostic assessments, instead relying on parental report. Given parent reported diagnoses of ASD have an estimated false-positive rate of 10-20% and false-negative rate of 5-25% (Warren et al., 2012), there are likely to be some individuals in the present study’s ASD group that are misclassified and an extremely small portion of the nASD and OPD group may actually have autism. However, the ADOS is not the only tool used to assess ASD (Grzadzinski et al., 2020) and the present study used the brief social responsiveness scale (SRS) to overcome the lack of formal diagnostic assessments. Individuals in the present study’s ASD group were removed from analysis if below previously established thresholds for reliable diagnosis of ASD using the SRS (Moul et al., 2015). This adjusted the initial number of individuals in the ASD group at baseline from 192 to 164.

The present study used a convenient sample from the ABCD study, focusing on a narrow period of adolescent development (9-12 years of age). Other investigations have reported neurodevelopmental changes across wide developmental ranges, but have required large sample sizes in order to do so (Khundrakpam et al., 2017). While the focus of the present study had the advantage of focusing on findings that may have otherwise been averaged out by deviations from other developmental periods, it does limit their generalization. The stability of reported findings at baseline and follow-up was also reported in post-hoc analysis in an attempt to ensure the stability of findings across short developmental periods was made transparent. Although this did help identify measures that were more robust across all time-points, it cannot account for changes from 12 years of age to adulthood or changes prior to 9 years of age. It is critical that future investigations focus on how these unique cytoarchitectural findings change across all of development.

## 5 Conclusion

Physiological associates to ASD have been historically difficult to reproduce due to the non-specific nature of macro-structural measures and inaccessibility of micro-structural measures. The current study uses a more recent methodology of measuring cytoarchitecture *in vivo* in a large population. While TND only identified a significant decrease in those with autism at the right cerebellum, TND was greater in those with autism diffusely throughout anterior fronto-temporal and basal ganglia regions, and TNI was decreased in the postero-occipital regions and hindbrain. Findings remained significant when verified in post-hoc comparison against other psychiatric disorders and at separate time points. Taken together, these findings demonstrate the unique importance of cytoarchitecture in further understanding the physiology of ASD and that the summation of RNI and RND into a single TND measure, may well obscure important findings related to the neuropathology of ASD. Future research is necessary to reproduce these findings and more clearly characterize how these heterogeneous findings interact at a global level.

## Author Contributions

ZPC performed the primary data analyses and statistical tests, created the illustrations, and wrote the first substantial draft of the manuscript. JJF and EGF are site PIs for the ABCD consortium at the University of Rochester and coordinated data collection and project management. ZPC, JJF and EGF collectively conceived of the study and consulted regularly during the development of the analyses and data representations. JJF and EGF provided editorial input to ZPC during multiple drafts of the manuscript.

All authors approve of this final version for publication, attest to the accuracy of the work reported, and agree to be fully accountable for all aspects of the work.

## Acknowledgements

Data used in the preparation of this article were obtained from the Adolescent Brain Cognitive Development^SM^ (ABCD) Study (https://abcdstudy.org), held in the NIMH Data Archive (NDA). This is a multisite, longitudinal study designed to recruit more than 10,000 children age 9-10 and follow them over 10 years into early adulthood. The ABCD Study® is supported by the National Institutes of Health and additional federal partners under award numbers U01DA041048, U01DA050989, U01DA051016, U01DA041022, U01DA051018, U01DA051037, U01DA050987, U01DA041174, U01DA041106, U01DA041117, U01DA041028, U01DA041134, U01DA050988, U01DA051039, U01DA041156, U01DA041025, U01DA041120, U01DA051038, U01DA041148, U01DA041093, U01DA041089, U24DA041123, U24DA041147. Neuroimaging at the University of Rochester ABCD site is conducted through the Translational Neuroimaging and Neurophysiology Core of the University of Rochester Intellectual and Developmental Disabilities Research Center (UR-IDDRC) which is supported by a center grant from the Eunice Kennedy Shriver National Institute of Child Health and Human Development (P50 HD103536 – to JJF). A full list of supporters is available at https://abcdstudy.org/federal-partners.html. A listing of participating sites and a complete listing of the study investigators can be found at https://abcdstudy.org/consortium_members/. ABCD consortium investigators designed and implemented the study and/or provided data but did not necessarily participate in the analysis or writing of this report. This manuscript reflects the views of the authors and may not reflect the opinions or views of the NIH or ABCD consortium investigators. The ABCD data repository grows and changes over time. The ABCD data used in this report came from <u>DOI: 10.15154/1523041</u>

